# Single-nuclear RNA sequencing reveals an ATF3-independent sensory-neuron program after surgical incision

**DOI:** 10.64898/2026.07.08.737350

**Authors:** Po-Yi Paul Su, Jessica Yu, Jarret Wenrich, Fang Ye, Lingyi Zhang, Zhonghui Guan

## Abstract

**Background:** Acute postoperative pain is common and often treated with opioids, but which sensory-neuron changes are responsible for the peripheral drive of pain is poorly defined. Skin incision induces activating transcription factor 3 (ATF3), the canonical marker of nerve injury, in dorsal root ganglion (DRG) neurons, and ATF3 marks the neurons that remain hyperexcitable. Whether ATF3 is required for postoperative pain is unknown.

**Methods:** Adult mice of both sexes underwent hind-paw plantar incision. Postoperative nociceptive behavior was compared between sensory-neuron–specific ATF3 conditional-knockout mice (*Avil*^Cre/+^; *Atf3*^fl/fl^) and littermate controls using assays that probe distinct primary-afferent modalities (von Frey; Hargreaves; dynamic light touch) together with an operator-independent index of spontaneous injury behavior. Single-nucleus RNA sequencing of wild-type and ATF3-null DRG sensory neurons (naïve and postoperative day 1) characterized the initiating injury program; transcription-factor–activity inference, RNAscope, and phospho–c-Jun immunostaining examined c-Jun; and the oral dual leucine zipper kinase (DLK) inhibitor GNE-3511, given at the time of incision, was tested behaviorally.

**Results:** Incision transiently induced ATF3 in a subset of DRG neurons, returning to baseline by day 14, in parallel with the two-week course of nociceptive behavior. Deleting ATF3 in sensory neurons did not change the onset or resolution of nociceptive behavior on any readout, in either sex, across mechanical, thermal, and tactile modalities. Single-nucleus profiling showed a broad surgery-responsive nociceptor program in which ATF3 was the most strongly induced gene, yet only 23% of surgery-responsive neurons expressed it. Without ATF3, the program was remodeled but not abolished, and transcription-factor–activity inference nominated c-Jun as a candidate ATF3-independent factor; incision induced c-Jun in vivo. Systemic inhibition of DLK, which activates c-Jun pathway, lowered c-Jun phosphorylation and reduced evoked nociceptive hypersensitivity.

**Conclusions:** Our results indicate that ATF3 marks part of the injured sensory neuron but does not drive acute post-surgical nociceptive behavior, and that it is not required for the postsurgical pain behaviors across afferent modalities carried by molecularly distinct nociceptor classes. The underlying injury program is broader than ATF3 and is nominated to depend on c-Jun. Because DLK inhibition, given at the time of incision, reduced evoked hypersensitivity, we propose the DLK–JNK–c-Jun axis as a candidate perioperative non-opioid target, which will require further genetic validation.

## 1. Introduction

Acute postoperative pain is one of the most common complications of surgery. In a national survey, about 80% of surgical patients reported postoperative pain, most of it moderate to extreme, and postoperative pain remains widely undertreated despite decades of attention; it prolongs recovery and drives continued perioperative opioid use.^1^ Incisional pain has long been attributed to inflammation, and it is modeled preclinically by hind paw plantar incision, which produces reliable and quantifiable hypersensitivity.^2^ However, what happens within the primary sensory neuron after surgery, and which of these changes actually drive pain, is still poorly defined.

Surgery is not a purely inflammatory insult to sensory neurons. Even without frank peripheral nerve transection, skin incision induces activating transcription factor 3 (ATF3) and other axonal-regeneration-associated genes in dorsal root ganglion (DRG) neurons, a molecular signature classically associated with nerve injury, likely because the nerve endings are cut during surgical incision.^3^ The neurons that carry this signature stay electrophysiologically hyperexcitable long after the wound has healed and inflammation has resolved, and ATF3 expression predicts which neurons remain sensitized.^4^ Consistent with this, plantar incision produces long-lasting intrinsic hyperexcitability in DRG pain sensing neuron (nociceptor) somata that persists for weeks.^5^ The transcriptional response to surgery, however, has been characterized only in bulk tissue, which averages across the molecularly distinct subtypes of sensory neurons and obscures them with other cell types within DRG.^6^ Cell-type-resolved analyses of injury-induced neuronal reprogramming, conducted by single-nuclear RNA sequencing (sn-RNA-Seq), have been applied to models of frank nerve injury,^7^ but the sensory-neuron response to surgery has not been resolved at the single-cell level, and whether surgery engages the subtype-spanning reprogramming program described after nerve injury is, to our knowledge, unknown.

ATF3 is absent in naïve sensory neurons and is rapidly and near-universally induced by axotomy, making it the canonical marker of peripheral nerve injury.^8^ Its established function, however, is regenerative. ATF3 raises the intrinsic growth state of DRG neurons and accelerates axon regeneration,^9^ and it is required for the broader reprogramming and functional recovery that follow nerve injury.^7^ ATF3 therefore marks the same neurons that become hyperexcitable after surgery, but with its known role to promote repair, it is unclear whether ATF3 drives the painful phenotype or is only a regenerative response that happens to coincide with it. The direct behavioral evidence whether ATF3 is involved in pain is limited and conflicting, and it comes entirely from non-surgical models: ATF3 knockdown does not alter nerve-injury-induced tactile allodynia,^10^ whereas ATF3 deletion prevents hyperalgesia in diabetic neuropathy.^11^ Nevertheless, whether ATF3 is required for postsurgical pain has not been tested.

Surgery is also unusual among painful conditions because the injury is planned and iatrogenic. The onset of noxious event has a known time and place, which opens a window to intervene at or just before incision; this is the premise of preemptive analgesia.^12^ We therefore focused on the initiation of nociceptive behavior over the first postoperative day, the phase in which such an intervention would act.

We addressed these two gaps directly. First, we defined the primary sensory-neuron response to hind paw incision at single-nucleus resolution; to our knowledge, this is the first single-nucleus characterization of the DRG in a plantar-incision surgical model. Second, using sensory-neuron–specific conditional ATF3 knockout mice, we tested whether ATF3 is required for the initiation and resolution of postoperative nociceptive behavior, using assays that probe distinct primary afferent modalities and an operator-independent index of spontaneous injury behavior. Single-nucleus profiling of ATF3-null and wild-type DRG then defined the ATF3-dependent component of that program in nociceptors. These experiments showed that the surgical injury program in nociceptors is broader than ATF3, and that the ATF3-independent nociceptor response is nominated to depend on the AP-1 transcription factor c-Jun. We then tested c-Jun’s upstream kinase, dual leucine zipper kinase (DLK), as a pharmacologic non-opioid target to reduce postsurgical pain behaviors.

## 2. Materials and Methods

### Animals

Approximately equal numbers of male and female young adult (8–12 weeks old) mice were used. Global *Atf3* knockout mice, maintained on a C57BL/6 background with no detectable developmental or behavioral abnormalities, were generously provided by Dr. Tsonwin Hai (Ohio State University);^13^ these and C57BL/6J wild-type mice (The Jackson Laboratory, or bred in-house from the same background) were used for single-nucleus RNA sequencing. The global knockout is a protein-null allele: the *Atf3* transcript is still produced, although cannot be translated into functional ATF3 protein, allowing injured neurons to be identified by the *Atf3* transcript in both genotypes (see Single-nucleus RNA sequencing). Behavioral experiments used sensory-neuron–specific conditional knockouts (*Avil*^Cre/+^; *Atf3*^fl/fl^), generated by crossing the Advillin-Cre driver^14^ to a floxed *Atf3* allele,^15^ together with their *Atf3*^fl/fl^ littermate controls. For paired ATF3-lineage/phospho–c-Jun immunofluorescence, *Atf3*-CreERT2^16^; *Rosa26*-YFP^17^ reporter mice were used, with ATF3 lineage read through the YFP reporter; tamoxifen (30 mg/mL in corn oil, 150 mg/kg intraperitoneally) was administered on postoperative day 1 to label the ATF3-induced (injured) lineage. Mice were housed under a 12-h light/dark cycle with food and water ad libitum, at up to 5 animals per cage; animals used for behavioral testing were never singly housed. Both sexes were studied; data were analyzed separately by sex and pooled when no sex difference was detected (see Statistical Analysis). All procedures were approved by the Institutional Animal Care and Use Committee of the University of California, San Francisco (protocol AN203447-00E). Euthanasia was performed by CO₂ inhalation followed by cervical dislocation.

### Hind paw incision

The hind paw incision model was performed as described previously (Pogatzki and Raja^18^). Briefly, under isoflurane anesthesia (2% isoflurane in 100% oxygen at 2 L/min) and aseptic conditions, a 5-mm longitudinal incision was made through the skin of the plantar surface of the left hind paw. The flexor digitorum brevis muscle was elevated and cut longitudinally, and the skin was closed with two 6-0 silk sutures. No postoperative analgesia was administered to avoid confounding the measurement of incision-induced nociceptive behavior. No sham-operated group was used; each animal served as its own control, compared with behavioral measures obtained at baseline before surgery.

### DLK inhibitor administration

The dual leucine zipper kinase (DLK) inhibitor GNE-3511^19^ was administered following the regimen of Wlaschin et al.,^20^ with modifications to timing. GNE-3511 was dissolved in 0.5% (w/v) USP-grade methylcellulose and 0.2% (v/v) Tween 80 in water, sonicated, and stored at 4 °C for no more than 7 days. Mice were dosed by oral gavage at 75 mg/kg (10 mL/kg of a 7.5 mg/mL suspension); control animals received an equal volume of vehicle. The first dose was given at the time of incision and repeated every 12 h for a total of three doses; the third dose (24 h after incision) preceded behavioral testing.

### Behavioral assessment

Behavioral experiments were performed between 9:00 am and 4:00 pm by an experimenter blinded to genotype and treatment. Animals were habituated to the apparatus for 3 h before baseline testing and for 1 h on each testing day. The sample size was based on previous experiments in our laboratory.^21^ Measurements were obtained at baseline (pre-incision) and on post-operative days 1, 3, 5, 7, 10, 14, and 21, as shown in the figures. The assays were chosen to sample distinct primary-afferent modalities.

#### Mechanical sensitivity (von Frey)

Punctate mechanical withdrawal thresholds were measured with calibrated von Frey filaments applied to the plantar surface of the hind paw. The 50% withdrawal threshold was determined by the Dixon up–down method^22^ with filaments from 0.04 to 2 g. A positive response was any paw lift or greater.

#### Thermal sensitivity (Hargreaves)

Radiant-heat thermal sensitivity was assessed with the Hargreaves test^23^ using the Ugo Basile Plantar Test apparatus (cat. #37570). Mice were placed on a glass surface, and a radiant heat beam (infrared intensity set to 30%) was focused on the plantar hind paw, with a 30-s cutoff to prevent tissue damage. Paw-withdrawal latency was recorded over 4 trials per session.

#### Dynamic light touch (light stroking)

Dynamic tactile sensitivity was assessed by light stroking, as a separate test from the Black Box assay. A soft paintbrush (5/0) was stroked across the plantar surface of the hind paw from heel to toe. The stimulus was applied three times at intervals of at least 5 min and scored on a 0–3 scale (0, no response; 3, sustained paw lifting/licking); the average score per animal was used.^24^

#### Automated gait analysis and the Spontaneous Injury Behavior Index

Gait and paw loading were measured on the Black Box 4-box platform (BlackBox Bio, Dallas, TX), which captures paw–floor contact by frustrated total internal reflection and derives paw luminance ratios (relative weight borne per paw) together with machine-learning body-pose estimation. Mice were recorded for 3–5 min per session and analyzed with the machine-learning behavioral-phenotyping pipeline of Layne et al.^25^ A composite, operator-independent Injury Behavior Index of spontaneous injury behavior was computed as the primary gait outcome, as the left-minus-right (injured-minus-uninjured) laterality of seven Black Box measures across three domains (paw-contact posture, gait timing, and weight bearing), each standardized to the group’s pre-incision baseline (z-score) and sign-oriented so that positive values indicate a more injury-like state; domains were combined hierarchically with equal weight (construction shown in Supplementary Fig S1). To control for possible sedative or motor effects, voluntary walking speed was extracted from the same recordings.

### Tissue collection and histology

For immunostaining, mice were deeply anesthetized, perfused transcardially with PBS, and tissues were post-fixed in 4% formalin for 3 h, cryoprotected in 30% sucrose, and embedded in OCT. For RNAscope in situ hybridization, mice were perfused transcardially with PBS as above, but the dorsal root ganglia were dissected fresh, flash-frozen on dry ice, and embedded in OCT without fixation. Lumbar dorsal root ganglia (L4 unless otherwise specified) were cryosectioned at 10 µm. (For the conditional-knockout validation only, lumbar spinal cord was also collected and sectioned at 30 µm to confirm sparing of axotomized ventral-horn motor neurons in Supplementary Fig S2; spinal cord was not otherwise part of this study.) For paired contralateral/ipsilateral comparisons, both sides were mounted on the same slide and processed and imaged together.

#### Immunohistochemistry and immunofluorescence

Cryosections were preincubated in blocking buffer (10% normal horse serum, 0.3% Triton X-100 in 0.1 M phosphate buffer), incubated with primary antibodies in blocking buffer, washed three times in 0.1 M PBS, and incubated with goat fluorescent secondary antibodies (Thermo Fisher; Alexa Fluor 488 #A-11008 or Alexa Fluor 555 #A-21428, as appropriate to the channel). The primary antibodies were: rabbit anti-ATF3 (Sigma-Aldrich #HPA001562, 1:500), used to detect ATF3 protein in DRG (Figure 1) and to confirm deletion (Supplementary Fig S2); rabbit anti–phospho–c-Jun (Ser63) (Cell Signaling Technology #9261S, 1:1000); and chicken anti-GFP (Abcam #ab300643, 1:2000) to detect the YFP reporter. Because the anti-ATF3 and anti–phospho–c-Jun antibodies are both raised in rabbit and cannot be co-stained, in the reporter experiments, ATF3 lineage was read through the YFP reporter (chicken anti-GFP with an Alexa Fluor 488 anti-chicken secondary, which is non–cross-reactive with the anti-rabbit secondary) in *Atf3*-CreERT2; *Rosa26*-YFP mice, while phospho–c-Jun was detected separately. Sections were counterstained with NeuroTrace 640/660 Deep-Red Fluorescent Nissl Stain (Invitrogen/Thermo Fisher #N21483) and DAPI. Images were acquired on an Olympus FV3000 laser-scanning confocal microscope (20× objective, z-stacks reduced to maximum-intensity projections), with identical laser power, gain, and exposure across compared images. Nuclei and neuronal somata were segmented with fine-tuned Cellpose/Omnipose models^26^ followed by manual curation. Per-neuron background-subtracted mean fluorescence intensity was measured, and a neuron was scored positive if it exceeded a defined threshold (for ATF3, a two-component Gaussian-mixture intersection; for phospho–c-Jun/YFP, the 95th percentile of that animal’s contralateral distribution). Images were analyzed in ImageJ (National Institutes of Health) and custom Python scripts.

**Figure 1.**
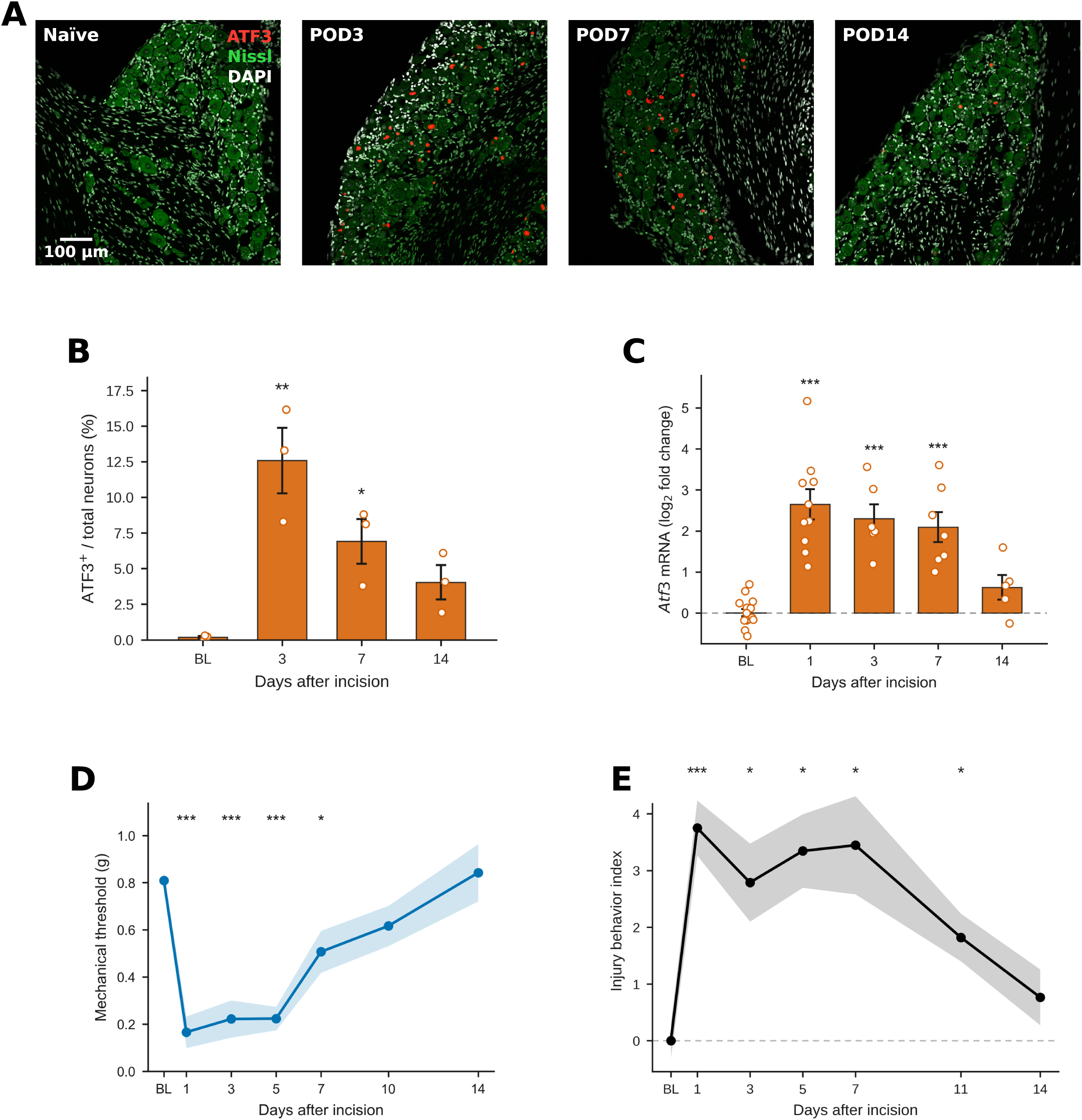
Hind-paw incision induces a transient ATF3⁺ sensory-neuron injury signature whose time course parallels nociceptive behavior. (A) Representative L4 dorsal root ganglion (DRG) immunostaining for ATF3 (red, nuclear), Nissl (green), and DAPI (white) in naïve mice and at postoperative day (POD) 3, 7, and 14 after plantar incision; ATF3 is undetectable in naïve DRG and induced in a subset of neurons after incision. Scale bar, 100 µm. (B) ATF3⁺ neurons (% of total; mean ± SEM, n = 3 per timepoint): uninjured 0.18 ± 0.09, POD3 12.58 ± 2.30, POD7 6.91 ± 1.57, POD14 4.03 ± 1.21; induction peaks at POD3 and resolves toward baseline by POD14 (one-way ANOVA P = 0.0026). (C) Atf3 mRNA (log₂ fold change) over the same axis, tracking the protein. (D) Mechanical withdrawal threshold in wild-type mice (von Frey, g); hypersensitivity develops and resolves in parallel with ATF3 (linear mixed-effects model, effect of time P < 0.0001). (E) Operator-independent injury-behavior index (n = 5). Mean ± SEM; *P < 0.05, **P < 0.01, ***P < 0.001.

#### RNAscope in situ hybridization

Multiplex fluorescent RNAscope (RNAscope Multiplex Fluorescent Reagent Kit v2; Advanced Cell Diagnostics [ACD]/Bio-Techne) was performed on 10-µm fresh-frozen DRG sections using probes against mouse *Atf3* (ACD #426891; C1, 488 nm) and *Jun* (ACD #453561; C3, 647 nm), with Nissl and DAPI counterstain. Contralateral and ipsilateral sections were mounted on the same slide and processed together. Images were acquired on an Olympus FV3000 confocal microscope (10× objective, single optical planes, pixel size 0.621 µm), with identical acquisition settings across images. Neurons were segmented with a fine-tuned Cellpose/Omnipose model and manually curated; per-neuron mean fluorescence intensity of *Atf3* and *Jun* was measured after background subtraction. *Atf3*-positive neurons were classified using a single global threshold (Gaussian-mixture intersection); *Jun*-high neurons were classified after per-animal contralateral normalization as those above the 75th percentile of the pooled normalized contralateral distribution.

### Single-nucleus RNA sequencing (snRNA-Seq)

#### Nuclei isolation, library preparation, and sequencing

Single-nucleus RNA sequencing was performed on dorsal root ganglia from wild-type (C57BL/6) and *Atf3* global-knockout mice under naïve conditions and one day after hind paw incision. Eight libraries were generated per genotype (wild-type: 3 naïve, 5 incision; Atf3-knockout: 4 naïve, 4 incision). Nuclei were isolated with the 10x Genomics Chromium Nuclei Isolation Kit; the L3 and L4 DRG were dissected and pooled, with 2–3 animals per library. Libraries were prepared with the 10x Genomics Chromium Single Cell 3′ v3 assay and sequenced on an Illumina NovaSeq X Plus at the UCSF Center for Advanced Technology, targeting 30,000–40,000 read pairs per nucleus (above the ≥20,000 read-pairs-per-nucleus minimum recommended by 10x Genomics for 3′ gene expression).

#### Alignment, quality control, and annotation

Reads were aligned with Cell Ranger 9.0.1 against GRCm39-2024-A with intron inclusion enabled. Because the Atf3-knockout allele is exon-specific and 3′ priming captures transcript fragments downstream of the deleted exon, the *Atf3* transcript is still detected in the knockout and was used to help identify injured neurons. Ambient RNA was removed per library with CellBender; nuclei with ≥500 genes and <10% mitochondrial reads were retained; doublets were removed with DoubletFinder^27^ and scDblFinder. Data were processed in Seurat v5,^28^ and functional sensory-neuron subtypes were annotated by label transfer from the harmonized DRG atlas of Bhuiyan et al.^29^ Analyses focused on the peptidergic (PEP; *Calca*⁺) and non-peptidergic (NP; *Mrgprd*⁺) nociceptor populations. Because wild-type and knockout libraries were sequenced in separate batches, genotypes were placed in a common space by reference projection (SCTransform, 30-PC PCA, stored UMAP; MapQuery; prediction-score floor 0.5).

#### Injury-responsive population, differential expression, and transcription-factor analysis

Injury-responsive nuclei were identified by neighborhood differential-abundance testing with Milo^30^ (k = 30, d = 30, spatial FDR < 0.10). Because incision markedly induces *Atf3*, a surgery-responsive union set was defined as nuclei that were Milo-enriched or expressed *Atf3* at ≥ 2 UMIs in the incision condition (the 2-UMI threshold kept the naïve false-positive rate below 0.3% in wild-type). Within each genotype, per-sample pseudobulk profiles (surgery-responsive incision vs. all naïve) were compared with edgeR^31^ quasi-likelihood F-tests and Benjamini– Hochberg correction. Eight literature-anchored functional gene sets were scored per nucleus with Seurat AddModuleScore, and per-replicate incision-minus-naïve deltas were derived. Transcription-factor target sets were taken from DoRothEA (A–C) regulons^32^ via decoupleR^33^ with a curated AP-1 regulon; regulon over-representation was tested by one-sided Fisher exact tests (Benjamini–Hochberg), and transcription-factor activity was estimated from log-fold-change vectors by univariate linear models (decoupleR).

### Statistical analysis

Data are presented as mean ± SEM, and P < 0.05 was considered significant. The experimental unit was the individual animal for behavioral and histological analyses, and the individual library (animal) for single-nucleus analyses. Behavioral inferential statistics were performed in R (rstatix); histology and single-nucleus statistics were computed in Python (scipy) and R (Seurat, miloR, edgeR, decoupleR) as above. A fixed random seed (42) was used for all stochastic steps. For behavioral time courses, a two-way mixed-model repeated-measures ANOVA was used (between: genotype or treatment; within: day), with Greenhouse–Geisser correction when sphericity was violated and generalized eta-squared as the effect size; analyses were complete-case, and per-timepoint post-hoc comparisons were not performed when the omnibus genotype and interaction terms were non-significant (ANOVA gatekeeping). For single-group injury time courses, a one-way repeated-measures ANOVA was used, with each post-incision timepoint compared to baseline by Holm–Bonferroni-corrected paired t-tests. Pooling across sex was justified by a linear mixed model (value ∼ genotype × sex × day + (1|animal)); a sex × day effect for thermal latency (a known baseline thermal sex difference; Sorge et al.^34^) did not involve genotype and did not preclude pooling. For histology, per-animal percentages of positive neurons were compared between sides by paired two-sided t-tests. For single-nucleus data, differential abundance used Milo (spatial FDR < 0.10), pseudobulk differential expression used edgeR with Benjamini–Hochberg correction, per-module cross-genotype deltas used two-sided Wilcoxon rank-sum tests (Benjamini–Hochberg), and regulon over-representation used one-sided Fisher exact tests (Benjamini–Hochberg).

### Data and code availability

Single-nucleus RNA sequencing data will be deposited in the NCBI Gene Expression Omnibus before publication; the accession number will be provided upon manuscript acceptance. The analysis code will be made publicly available in a version-controlled repository upon manuscript acceptance. Behavioral, gait, histology, and source data are available from the corresponding author upon reasonable request.

## 3. Results

### Surgical incision induces a transient ATF3⁺ induction in DRG neurons with similar time course of postsurgical pain behavior

Because ATF3 is the classical canonical marker of injured DRG neurons, we first examined its expression over the time course of a plantar incision. ATF3 protein was undetectable in naïve L4 dorsal root ganglion (DRG) and was induced in a discrete subset of neurons after unilateral plantar incision. The percentage of ATF3⁺ neurons rose from 0.18 ± 0.09% at baseline to 12.58 ± 2.30% on postoperative day (POD) 3 and returned toward baseline by POD14 (4.03 ± 1.21%; one-way ANOVA *P* = 0.0026; Fig 1A, 1B). Notably, the *Atf3* mRNA induction followed the same time course (Fig 1C).

The postsurgical pain behavioral time course matched the time course of ATF3 induction. Mechanical withdrawal thresholds fell after incision and recovered over the same two weeks (Fig 1D), as did assessment of operator-independent spontaneous injury behavior from the Black Box platform (Fig 1E; index construction in Supplementary Fig S1). ATF3 induction and resolution, therefore, coincided with the onset and recovery of nociceptive behavior.

### Sensory-neuron ATF3 is not required for acute post-surgical nociceptive behavior across afferent modalities

We have deleted *Atf3* gene selectively in sensory neurons (*Avil*^Cre/+^; *Atf3*^fl/fl^) and compared incisional nociceptive behavior between *Avil*^Cre/+^; *Atf3*^fl/fl^ mice and +/+; *Atf3*^fl/fl^ littermate controls over 14 days. The behavioral battery was chosen to probe distinct primary-afferent modalities rather than a single “pain” readout: punctate mechanical withdrawal (von Frey), which depends on non-peptidergic MrgprD⁺ C-fibers and Aδ mechanonociceptors;^35^ radiant heat (Hargreaves), which depends predominantly on TRPV1⁺ peptidergic nociceptors;^35^ and dynamic light touch (brush), which is signaled by Aβ low-threshold mechanoreceptors.^24^ We also measured spontaneous injury behavior. Despite the robust ATF3 induction shown in Figure 1, *Atf3* deletion from sensory neurons did not change the magnitude or the time course of any of these measures. Onset and resolution were indistinguishable between genotypes for mechanical threshold, thermal latency, dynamic light touch, and the spontaneous injury index, with no genotype main effect (von Frey *P* = 0.13; Hargreaves *P* = 0.25; light touch *P* = 0.31; injury index *P* = 0.47) and no genotype × time interaction (all *P* ≥ 0.27), whereas incision produced a strong main effect of day on every measure (all *P* < 0.0001; Fig 2). ATF3 is therefore not required across afferent modalities carried by molecularly distinct nociceptor classes, including the peptidergic and non-peptidergic populations profiled below.

**Figure 2.**
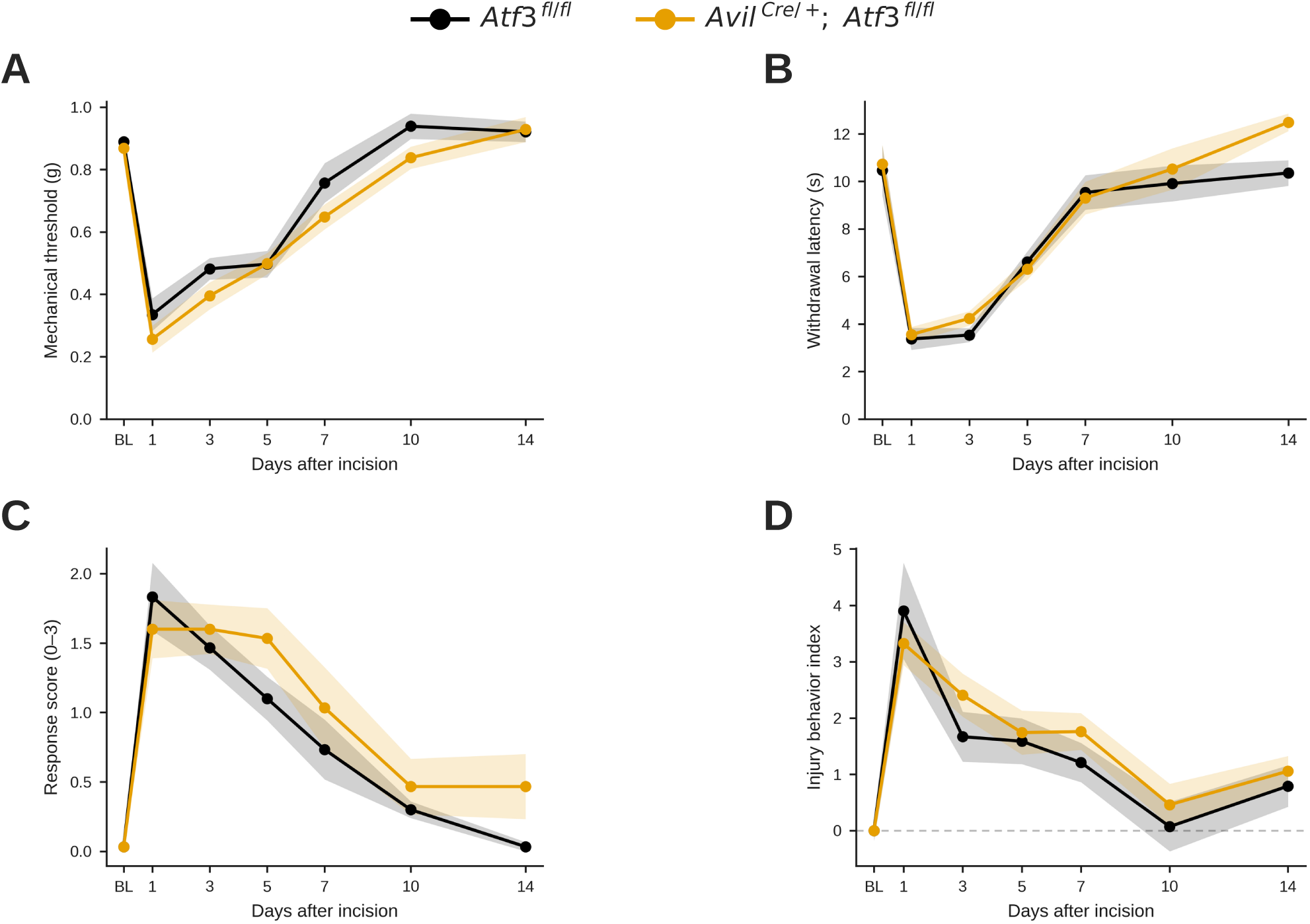
Sensory-neuron ATF3 deletion does not alter the initiation or recovery of acute post-surgical nociceptive behavior. Behavior after plantar incision in *Avil*^Cre/+^; *Atf3*^fl/fl^ conditional-knockout mice versus *Atf3*^fl/fl^ littermate controls: (A) mechanical withdrawal threshold (von Frey, g), (B) thermal latency (Hargreaves, s), (C) dynamic light-touch score, and (D) injury-behavior index over 14 days (n = 10 per genotype for A–C [5 female, 5 male]; n = 11 for D [6 female, 5 male]). Onset and resolution are indistinguishable between genotypes, with no genotype main effect (von Frey P = 0.13; Hargreaves P = 0.25; light touch P = 0.31; injury index P = 0.47) and no genotype × time interaction (all P ≥ 0.27; mixed-model repeated-measures ANOVA), whereas incision drove a robust main effect of day on every readout (all P < 0.0001). Protein-level knockout validation is shown in Supplementary Figure S2 (ATF3⁺ neurons present in wild-type DRG, absent in global Atf3-null and conditional-knockout DRG, and retained in conditional-knockout spinal motor neurons, confirming a sensory-restricted deletion). Mean ± SEM; n.s., not significant.

This negative result held up under several checks. It was reproduced within each sex, and voluntary walking speed was genotype-independent, which argues against a generalized motor confound (Supplementary Fig S3). Immunostaining confirmed the deletion was sensory-restricted: axotomized DRG neurons lacked ATF3 protein in the conditional knockout, whereas axotomized spinal motor neurons in the same animals retained ATF3 induction (Supplementary Fig S2). These results indicate that incision induces ATF3 but does not require it for acute post-surgical nociceptive behavior, and that the ATF3⁺ signature marks the injured neuron rather than drives its painful phenotype.

### Single-nucleus RNA sequencing resolves a broad surgery-responsive nociceptor program

Because ATF3 marks injured neurons but does not shape nociceptive behavior, we next asked what larger program is initiated in nociceptors after incision. We profiled 22,736 wild-type DRG neurons (3 naïve, 5 POD1) by snRNA-Seq and annotated eight canonical neuronal subtypes (Fig 3A; quality control in Supplementary Fig S4). We first focused on the peptidergic nociceptors (PEP_Calca). Neighborhood differential-abundance testing identified a surgery-responsive (SR) population, as described in the Material and Method section, that emerged within this lineage after incision; the SR fraction rose from 0.24 ± 0.24% in naïve tissue to 14.14 ± 1.82% at POD1 (Mann–Whitney *P* = 0.036; Fig 3B).

**Figure 3.**
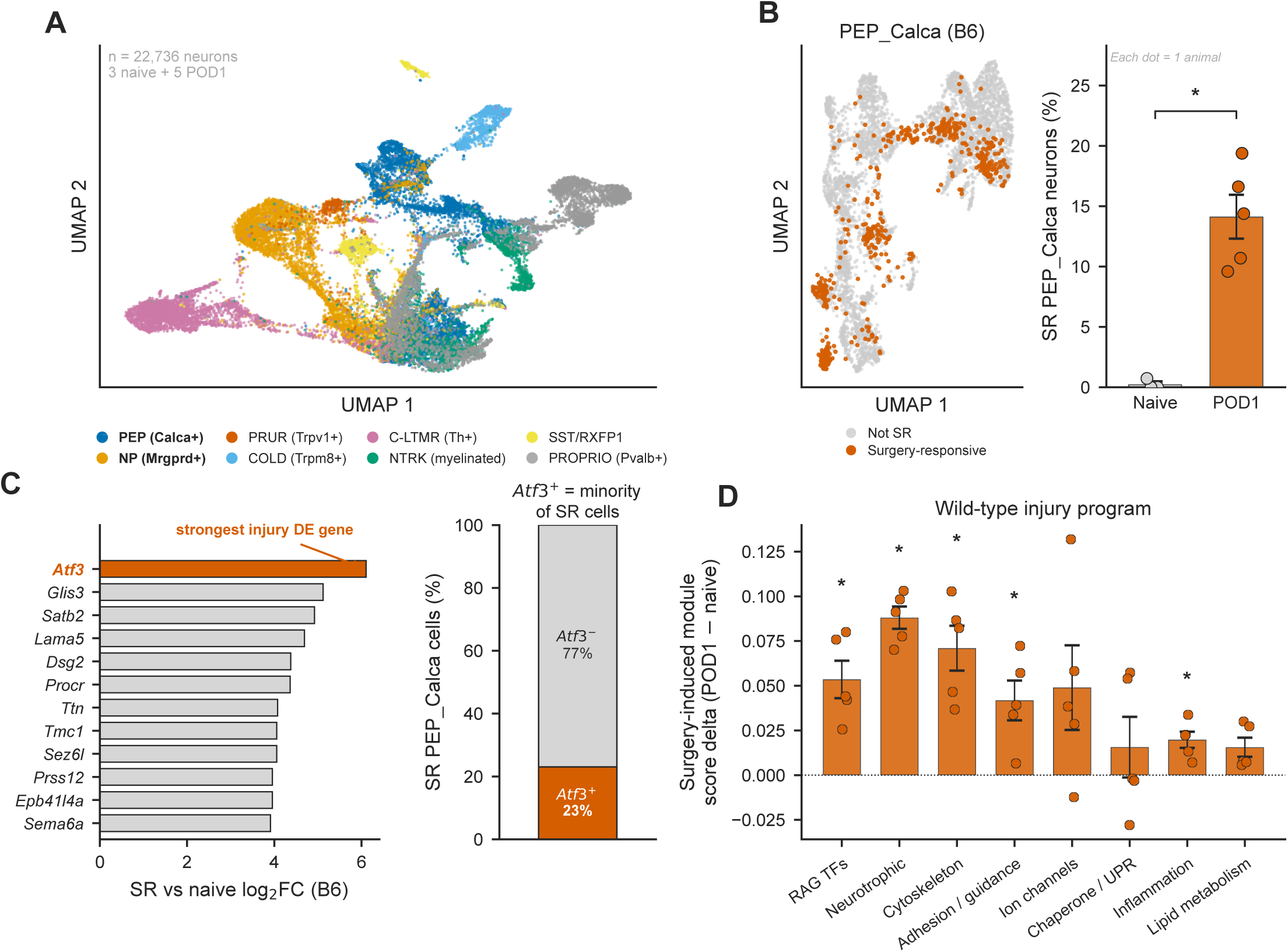
Single-nucleus RNA sequencing resolves a defined surgery-responsive program in peptidergic nociceptors. (A) UMAP of 22,736 wild-type DRG neurons (3 naïve, 5 POD1), annotated into eight canonical subtypes; peptidergic (PEP_Calca) and non-peptidergic (NP_Mrgprd) nociceptors highlighted. (B) A surgery-responsive (SR) population within peptidergic nociceptors, identified by neighborhood differential-abundance testing (Milo; spatial FDR < 0.10) with an Atf3-induction criterion; the per-animal SR fraction rises from 0.24 ± 0.24% (naïve) to 14.14 ± 1.82% (POD1) (mean ± SEM; n = 3 naïve, 5 POD1; two-sided Mann–Whitney P = 0.036). (C) The response is broader than the canonical marker: Atf3 is the most strongly induced gene (log₂FC 6.1; FDR 3 × 10⁻^6^ among 911 induced genes), yet only 23% of SR nuclei express detectable Atf3. (D) Injury-induced module scores (POD1 − naïve, mean ± SEM per replicate) across eight functional categories; regeneration-associated-transcription-factor, neurotrophic, cytoskeleton, adhesion/guidance, and inflammation modules are induced (two-sided Mann–Whitney P = 0.036 each), while ion-channel, chaperone/UPR, and lipid modules trend up without reaching significance at this replicate depth. Mean ± SEM; *P < 0.05.

The surgery incision-induced response of sensory neurons was broader than typical neuronal injury marker. Atf3 was the most strongly induced gene in the surgery-responsive population (log₂FC 6.1; FDR 3 × 10⁻^6^; rank 1 of 911 induced genes), yet only 23% of SR nuclei carried detectable Atf3 (Fig 3C). Most surgery-responsive neurons therefore executed the program without expressing the classical marker of Atf3; part of this gap may reflect incomplete transcript capture by single-nucleus sequencing rather than true absence of Atf3, given the higher concordance of the Jun and Atf3 signals between snRNA-seq and RNAscope/immunostaining (below). Scoring the program across eight literature-curated functional categories seeded from the genes up-regulated in the SR population, we found the regeneration-associated transcription-factor, neurotrophic, cytoskeletal, adhesion/axon-guidance, and inflammation modules coordinately induced (Mann–Whitney P = 0.036 each), while the ion-channel, chaperone/UPR, and lipid modules trended upward without reaching significance at this replicate depth (Fig 3D).

### Without ATF3, specific program modules are altered, and the residual response is nominated to depend on c-Jun

We repeated the profiling in Atf3-knockout DRG and compared the injury response by genotype (quality control, Supplementary Fig S5; naïve genotypes were transcriptionally equivalent, which excludes a developmental confound, Supplementary Fig S6). Loss of ATF3 did not abolish the injury response. The SR fraction was comparable between genotypes (14.1% vs 18.2%; Mann–Whitney P = 0.19), so what changed was the composition of the program, not its size (Fig 4A). Category by category, the structural adhesion/axon-guidance and ion-channel modules were attenuated in the knockout (module-score delta +0.042 → −0.007, P = 0.016; +0.049 → −0.017, P = 0.032), whereas the chaperone/UPR and lipid modules were increased, and the regeneration-associated-TF, cytoskeletal, neurotrophic, and inflammation modules were unchanged (Fig 4B). No functional category survived Benjamini–Hochberg correction across the eight tests; we therefore report raw P values from an underpowered comparison and describe an ATF3-dependent remodeling of which modules are engaged, chiefly the structural and ion-channel modules, rather than a wholesale change in the program.

**Figure 4.**
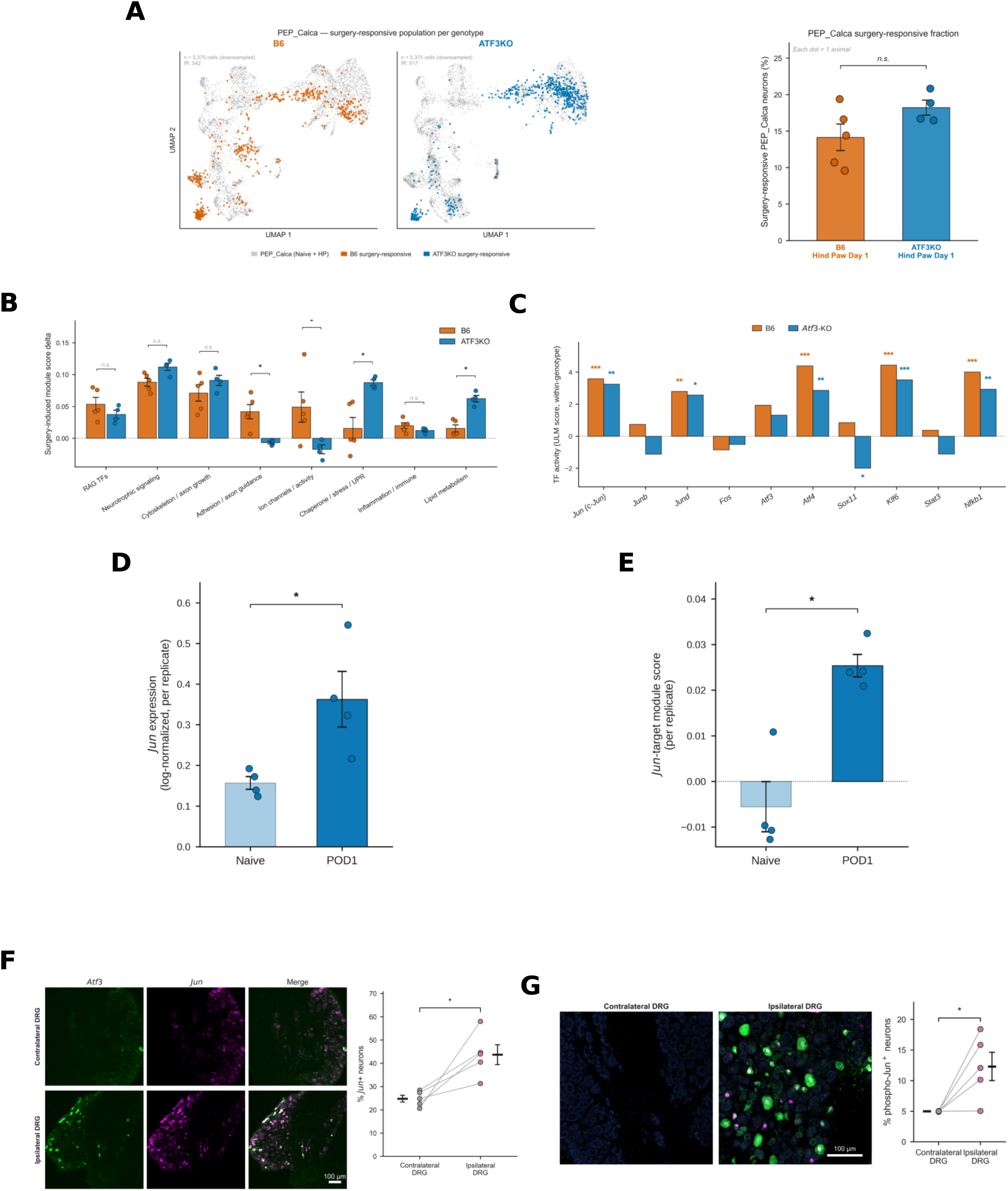
In the absence of ATF3, the injury program is remodeled, and its residual output is nominated to depend on c-Jun, which is induced by incision in vivo. (A) The surgery-responsive (SR) population within peptidergic (PEP_Calca) nociceptors on the per-genotype UMAP (wild-type and Atf3-knockout SR nuclei highlighted); the SR fraction is comparable between genotypes (14.1% vs 18.2%; two-sided Mann–Whitney P = 0.19, n.s.), so ATF3 loss changes program composition, not the size of the response. (B) Functional-category engagement by genotype: in Atf3-knockout, the adhesion/axon-guidance and ion-channel categories are attenuated (module-score delta +0.042 → −0.007, P = 0.016; +0.049 → −0.017, P = 0.032), whereas chaperone/UPR and lipid categories are preserved or increased, and regeneration-associated-transcription-factor, cytoskeleton, neurotrophic, and inflammation categories are unchanged. Two-sided Wilcoxon on per-replicate deltas (wild-type n = 5, knockout n = 4); no category survives Benjamini–Hochberg correction across the eight tests, and the panel reports raw P. (C) Transcription-factor activity inference (decoupleR univariate linear model) nominates c-Jun: Jun-family activity is preserved across genotypes (Jun 3.58 → 3.25; Jund 2.80 → 2.56). (D) Jun transcript is induced within the Atf3-knockout (POD1 versus naïve, P = 0.029). (E) The injury-induced increase in the Jun-target module footprint is significant within the Atf3-knockout (POD1 vs naïve, P = 0.029); because the absolute Jun⁺ fraction is lower in the knockout and this readout carries UMI-detection sensitivity, the comparison is interpreted cautiously. (F) Dual RNAscope for Jun and Atf3 (L4 DRG, POD1) shows Jun induced ipsilaterally in both Atf3⁻ and Atf3⁺ neurons (24.8% → 43.7% Jun⁺; paired two-sided t-test, P = 0.022, n = 5). (G) Phospho–c-Jun immunostaining (ipsilateral vs contralateral, paired per animal; 5.0% → 12.3%; paired two-sided t-test, P = 0.034, n = 5) shows JNK–c-Jun activation in tissue. Scale bars (F, G), 100 µm. Mean ± SEM; n.s., not significant.

To identify what sustains the response without ATF3, we used transcription-factor activity inference, which nominated several regeneration-associated factors. We focused on c-Jun for two reasons: its activity was preserved across genotypes, and prior single-cell reprogramming work independently identifies c-Jun as a principal transcription factor of the injured sensory neuron.^7^ Jun-family activity was preserved across genotypes (Jun 3.58 → 3.25; Jund 2.80 → 2.56), and an orthogonal regulon-overlap analysis supported the nomination at the target-set level, with c-Jun targets most over-represented in the neurotrophic and chaperone/UPR modules (Supplementary Fig S7). Within the knockout, both Jun transcript (Fig 4D) and the Jun-target module footprint (Fig 4E) were induced by incision (P = 0.029), and the module increase was greater in the knockout than in wild-type (between-genotype delta, Mann–Whitney P = 0.016). The same Jun-induction pattern appeared in the mechanistically distinct non-peptidergic (NP_Mrgprd) lineage in an underpowered sensitivity check, consistent with generality across nociceptor classes, though not establishing it (Supplementary Fig S8).

We corroborated the snRNA-seq Jun-induction result in tissue. Dual RNAscope for Jun and Atf3 showed Jun was induced in the ipsilateral DRG at POD1 (24.8% → 43.7% Jun⁺; paired t-test P = 0.022, n = 5), in both Atf3⁻ and Atf3⁺ neurons, consistent with an ATF3-independent source (Fig 4F). Phospho–c-Jun immunostaining showed activation of the pathway at the protein level (5.0% → 12.3% p–c-Jun⁺; paired t-test P = 0.034, n = 5; Fig 4G). The survey, therefore, nominates c-Jun as a candidate ATF3-independent factor associated with the residual program, and incision induces c-Jun, including its phosphorylated active form, in vivo. Whether c-Jun is causally required for post-surgical nociceptive behavior requires a specific neuronal c-Jun deletion or modulation and is the subject of a future conditional-knockout study.

### Pharmacologic DLK inhibition lowers c-Jun phosphorylation and attenuates evoked post-surgical hypersensitivity

Finally, we tested the axis as a potential target to manage postsurgical pain. DLK, an axonal-injury kinase upstream of JNK, phosphorylates c-Jun. We administered the DLK inhibitor GNE-3511 orally, starting at the time of incision. GNE-3511 reduced the fraction of p–c-Jun⁺ DRG neurons at POD1 (5.1% → 1.2%; P = 0.023; Fig 5A), preventing the injury-induced c-Jun phosphorylation seen in vehicle-treated mice and confirming target engagement. GNE-3511 also reduced mechanical and thermal hypersensitivity at POD1 (von Frey P = 0.046; Hargreaves P = 1.1 × 10⁻^5^), in both sexes (Fig 5B). Walking speed was comparable across groups, arguing against a sedative confound.

**Figure 5.**
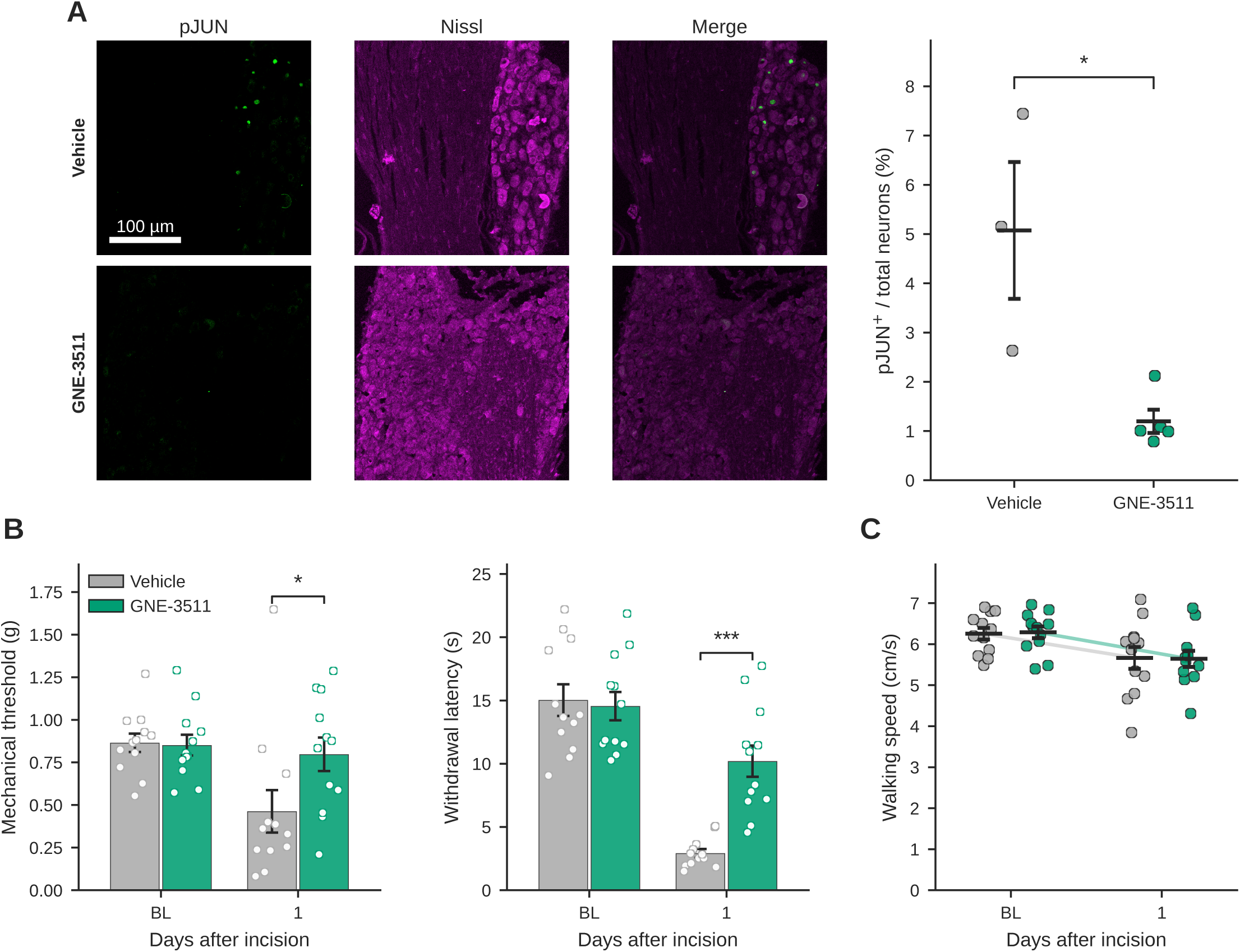
Pharmacologic DLK inhibition reduces c-Jun phosphorylation and attenuates evoked post-surgical hypersensitivity. (A) Oral GNE-3511 (DLK inhibitor) versus vehicle, given at the time of plantar incision; phospho–c-Jun immunostaining in L4 DRG at POD1. GNE-3511 reduces the fraction of p–c-Jun⁺ neurons (5.1% → 1.2%; log-transformed Welch t, P = 0.023; n = 3 and 5), confirming that DLK inhibition prevents the injury-induced c-Jun phosphorylation. (B) GNE-3511 attenuates mechanical (von Frey, P = 0.046) and thermal (Hargreaves, P = 1.1 × 10⁻^5^) hypersensitivity at POD1, in both sexes. (C) Walking speed declines comparably in all groups, so the reduced hypersensitivity is not attributable to drug-related sedation. Mean ± SEM.

Because ATF3 deletion alone does not alter nociceptive behavior (Fig 2) and c-Jun induction is ATF3-independent (Fig 4), the reduction in nociceptive hypersensitivity is most consistent with action through the ATF3-independent c-Jun axis.

## 4. Discussion

In this study, hind-paw incision reproduced the canonical ATF3⁺ nerve-injury signature in a subset of neurons within the lumbar DRG, and the induction and resolution of that signature tracked the onset and recovery of nociceptive behavior. However, deleting ATF3 in sensory neurons did not change the magnitude or the time course of postoperative nociceptive behaviors across both sexes and across afferent modalities carried by molecularly distinct nociceptor classes. Single-nucleus profiling showed how this dissociation is possible. Although ATF3 is the most strongly induced gene in the surgery-responsive nociceptor, it marks only a fraction of the neurons that execute a much broader, subtype-spanning injury program, and when ATF3 is absent, this program is remodeled rather than abolished. Transcription-factor activity inference nominates the AP-1 factor c-Jun as the candidate ATF3-independent driver of the residual output, and incision induces c-Jun in vivo at both the transcript and the phospho-protein level; this is induction, not demonstrated necessity. Systemic inhibition of DLK, the injury kinase upstream of c-Jun, lowered c-Jun phosphorylation and reduced evoked postoperative hypersensitivity. Because the inhibitor is systemically administered and DLK also acts in non-neuronal cells, these effects are associated with, but not mechanistically pinned to, neuronal c-Jun. Together, these results indicate that ATF3 is a marker of the injured neuron rather than a driver of post-surgical pain, and they nominate an ATF3-independent DLK–JNK–c-Jun axis as a candidate point of therapeutic intervention.

Our finding that ATF3 is not required for post-surgical pain resolves an apparent paradox in the literature. ATF3 is the classical marker of peripheral nerve injury,^8^ and it is also a genuine driver of the neuron’s regenerative response: it raises the intrinsic growth state of dorsal root ganglion (DRG) neurons^9^ and is required for the subtype-spanning reprogramming and functional recovery that follow axotomy.^7^ Because incision induces ATF3 in the same neurons that remain hyperexcitable,^4^ it was reasonable to suspect the contribution of ATF3 in the painful phenotype. However, the behavioral genetics argue otherwise. They agree with the only prior loss-of-function data in a non-diabetic model, in which ATF3 knockdown did not alter nerve-injury induced allodynia;^10^ the opposing result in diabetic neuropathy^11^ may reflect a disease context in which the regenerative and metabolic functions of ATF3 act on a chronic metabolic insult rather than an acute injury. The simplest interpretation is that ATF3 governs how the injured neuron repairs itself, not whether it signals pain. This distinction matters because ATF3 is commonly used as a surrogate for the painful neuron, and the two have been conflated.

Resolving the response at the single-nucleus level clarified what ATF3 actually marks. Prior transcriptomic work on surgery was done in bulk tissue,^6^ and cell-type-resolved reprogramming had been described only after frank nerve injury.^7^ We found that surgical incision, even though it leaves the nerve in continuity, engages a coordinated program across a defined surgery-responsive nociceptor population, yet only about a quarter of those surgery-responsive neurons express detectable *Atf3* (a gap that partly reflects the limited transcript capture of single-nucleus sequencing). The canonical marker, therefore, underestimates the surgery-responsive population, which helps explain how a marker can be induced and can track nociceptive behavior without being required for it. When ATF3 was deleted, transcription-factor activity inference and in situ validation converged on c-Jun as the leading candidate factor associated with the residual output of the program. This fits a long line of regeneration biology in which c-Jun is a master AP-1 regulator of the injured neuron. c-Jun is induced by axotomy, is required for efficient axonal regeneration and target reinnervation,^36^ and cooperates with ATF3, including as ATF3–Jun heterodimers that promote neurite outgrowth,^37^ so that the two factors normally act together. Our activity-inference data are consistent with c-Jun carrying the residual program when ATF3 is absent, though this regeneration biology makes the possibility plausible rather than establishing it here. Being able to separate the pain-relevant arm of the injury response from regeneration would itself be valuable: an ideal perioperative target would blunt the drive to pain while leaving the neuron’s regenerative recovery intact, rather than slowing repair.

Placing c-Jun upstream of pain also connects our findings to the JNK–c-Jun literature and to a conserved neuronal injury module. JNK–c-Jun signaling has an established role in injury-evoked hypersensitivity, although prior work localized much of it to spinal astrocytes and to transient activation in DRG neurons after nerve ligation;^38^ our results point instead to a surgical incision-evoked neuronal arm. The kinase we targeted, DLK, is the apical sensor of this module. It is required for mechanical allodynia and microgliosis after nerve injury,^20^ and it couples axonal injury to the c-Jun transcriptional program that spans apoptotic and regenerative responses.^39^ The same DLK–JNK–c-Jun axis drives apoptotic and neurodegenerative responses in other neurons, including dopaminergic neuron degeneration.^40^ Post-surgical nociceptor reprogramming is therefore one context-specific output of a widely used stress-response pathway. This breadth is both the appeal of the target and a reason for caution, because the pathway is not specific to pain.

Our study makes three main advances. First, to our knowledge, it provides the first single-nucleus map of the DRG sensory neurons response to surgical hind-paw plantar incision, and shows that surgical incision engages a subtype-spanning reprogramming program that had been described predominantly after nerve transection. This reinforces that surgical incision is not a purely inflammatory event: it drives nerve-injury-like neuronal reprogramming in the DRG, which fits the clinical observation that operations involving nerve transection or traction carry a higher risk of chronic post-surgical pain.^41^ Second, it separates a canonical injury marker from the pain it accompanies: using conditional and global genetics, we show that ATF3 is induced in the sensory neurons after surgical incision but not required for postsurgical pain behaviors across mechanical, thermal, and tactile modalities. Third, it nominates an ATF3-independent c-Jun axis as a candidate effector of the residual program, and shows that systemic inhibition of the upstream kinase, given at the time of incision, reduces evoked hypersensitivity.

Several strengths support these conclusions. The negative ATF3 result rests on conditional genetic models, protein-level confirmation of the deletion, independent behavioral readouts that sample distinct primary-afferent modalities, and replication in both sexes. The single-nucleus comparison used a matched cross-genotype design with demonstrated baseline transcriptional equivalence. We also corroborated the c-Jun nomination in tissue, by dual in situ hybridization and by phospho–c-Jun immunostaining, which moves the claim from computation to protein-level evidence in vivo.

Our study has several limitations. First, the c-Jun result is a nomination. The cross-genotype transcriptomic analyses are underpowered; no functional category survived multiple-comparison correction, and the in vivo data show that c-Jun is induced, not that it is required, so its necessity for post-surgical pain remains unproven. Second, the pharmacologic test has its own caveats. GNE-3511 was given systemically,^19^ and DLK also acts in non-neuronal cells such as microglia,^20^ so we cannot attribute the analgesia to neuronal c-Jun specifically, and the behavioral effect was limited to evoked hypersensitivity rather than the spontaneous injury index. Finally, the single-nucleus survey captured only the initiation phase at postoperative day 1 in a rodent plantar-incision model, and it does not address the later resolution phase of pain. The decisive next experiment is genetic and cell-type-specific by generating a sensory-neuron-conditional *c-Jun* knockout to test whether neuronal c-Jun is necessary for the initiation and resolution of post-surgical pain, and to separate necessity from the sufficiency implied by our activity inference. A cell-type-restricted DLK deletion would similarly separate neuronal from non-neuronal contributions to the analgesia we observed with the inhibitor. It will also be important to define the c-Jun target genes that alter nociceptor excitability and to link them to the long-lasting intrinsic hyperexcitability that outlasts behavior after incision,^5^ which would connect the transcriptional program to a physiological mechanism. Extending the single-nucleus time course beyond the first postoperative day, and testing whether transient perioperative blockade of the axis changes the course toward persistent post-surgical pain,^42^ are the translationally important next steps.

These findings have a specific translational logic that follows from the nature of postsurgical pain. Surgery is a planned tissue injury. Unlike most painful conditions, the noxious event has a medically planned time and place, which creates a preemptive window to intervene before the neuron commits to its injury program.^12^ Current perioperative practice can interrupt nociceptive transmission through local and regional anesthetics and multimodal analgesia,^43^ but blocking conduction does not prevent the transcriptional and translational reprogramming that incision sets in motion, so the molecular injury response proceeds beneath even a well-placed block and can seed the peripheral and central sensitization that outlasts the anesthetic.^44^ A therapy that acts at the level of this reprogramming would complement, rather than duplicate, conduction blockade. Blocking the pathway at the time of incision, before sensitization is established, is the preemptive strategy that the biology suggests, with the goal of preventing the downstream sensitization that produces persistent post-surgical pain.^42^ The DRG is an attractive site for such an intervention, because it is anatomically accessible to anesthesiologists and pain physicians through transforaminal epidural and intrathecal routes that are already in clinical use. A molecularly targeted agent could therefore be delivered where the reprogramming occurs, while limiting systemic exposure. This last point matters: systemic DLK inhibition was poorly tolerated and produced dose-limiting neurological toxicity in an amyotrophic lateral sclerosis trial,^45^ which argues against chronic global dosing and in favor of a local, time-limited, perioperative use of the concept. Our data motivate, but do not test, the hypothesis that transient, DRG-directed inhibition of the DLK–JNK–c-Jun axis during the preemptive and immediate windows could reduce persistent post-surgical pain without impairing the neuron’s regenerative repair, in which c-Jun also participates.^36^ Testing this will require a chronic pain endpoint, a cell-type-restricted manipulation, and a targeted-delivery study, the goal of the future studies.

In summary, surgical incision induces the canonical ATF3⁺ injury signature in sensory neurons, but ATF3 itself is not required for acute post-surgical nociceptive behaviors across afferent modalities. The underlying neuronal reprogramming is broader than ATF3, and in the absence of ATF3, it is nominated to depend on an ATF3-independent c-Jun axis; inhibiting the upstream kinase of this axis at the time of incision reduced evoked hypersensitivity. Our results reposition a canonical injury marker as a bystander to the pain it accompanies, and nominate the DLK–JNK–c-Jun axis as a candidate therapeutic target for DRG-directed non-opioid intervention.

## Supplemental Digital Content

Supplementary Figures S1–S8 are provided as Supplemental Digital Content and are cited in the text. Significance: **\*P < 0.05,** **P < 0.01, ***P < 0.001; n.s., not significant. Values are mean ± SEM unless noted.

**Supplementary Figure S1.**
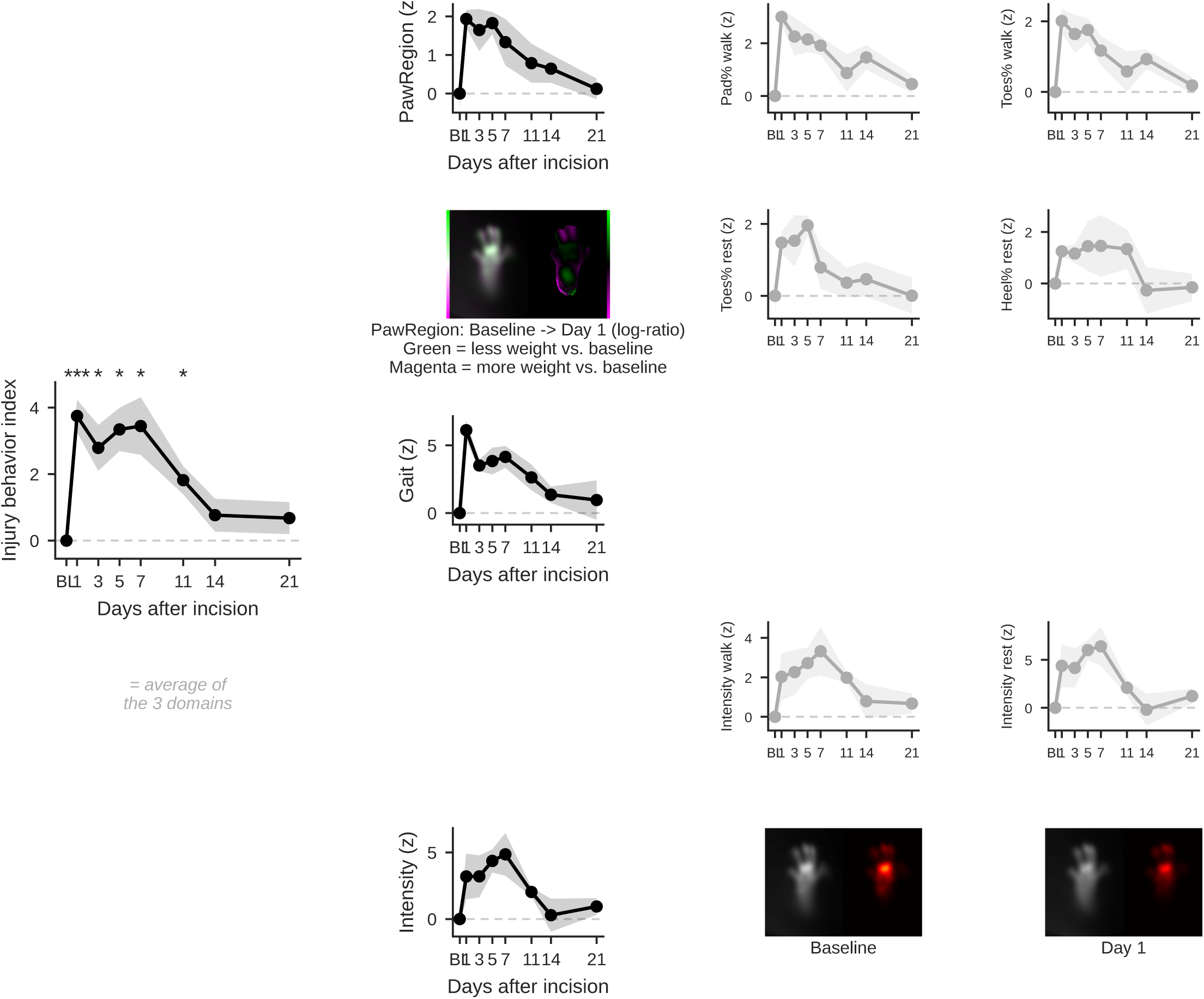
Derivation of the operator-independent Black Box injury-behavior index. Decomposition of the composite injury-behavior index used in Figures 1, 2, and 5, from a representative wild-type (C57BL/6) male cohort. The index is the left-minus-right (injured-minus-uninjured) asymmetry of hind-paw Black Box measures, standardized to each group’s pre-incision baseline (z-score, 0 = baseline) and sign-oriented so positive values denote more injury-like behavior; it combines three equally weighted domains (paw-contact distribution [posture], gait timing, and weight bearing [intensity]). The composite index rises sharply after incision and resolves toward baseline (one-way repeated-measures ANOVA across time; post-hoc paired t versus baseline with Holm–Bonferroni correction). The constituent domain z-scores and their component sub-measures (pad % and toes % during walking; toes % and heel % while stationary; step time; mean contact intensity during walking and while stationary) are shown for descriptive decomposition only and were not separately tested. Representative Black Box readouts are shown: the paw-region contact log-ratio (baseline → POD1; green = reduced and magenta = increased contact relative to baseline) and the weight-bearing (contact-intensity) images of both hind paws at baseline versus POD1. Day 8 was not analyzed (redundant with POD7); stride length was evaluated but excluded for failing to track the injury response.

**Supplementary Figure S2.**
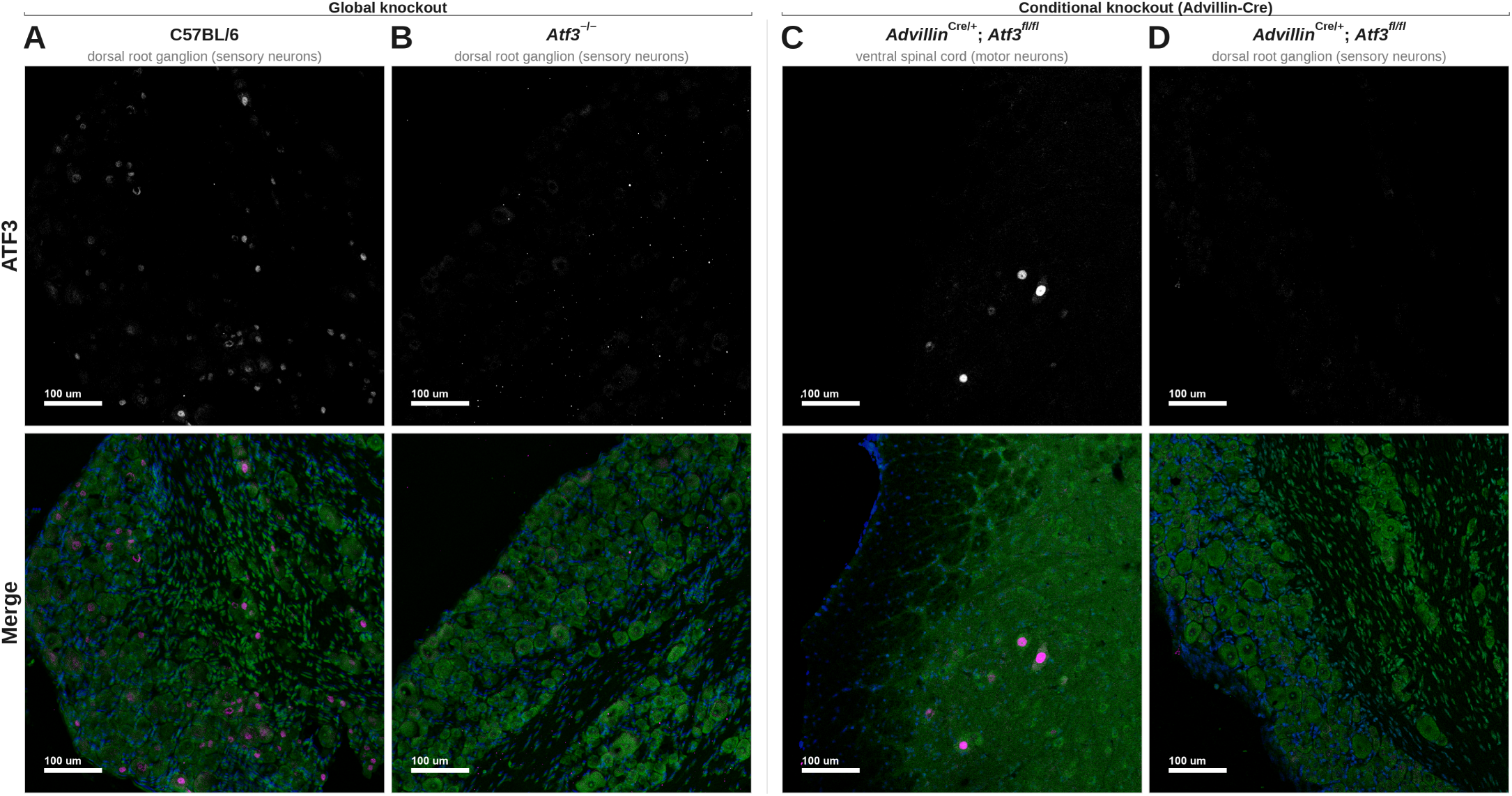
ATF3 protein is deleted in two Atf3 loss-of-function models (immunohistochemical validation). Support for Figure 2. Qualitative confirmation that the ATF3 protein is absent in the Atf3 loss-of-function models, corroborating the genotyping. Representative confocal images of DRG and spinal cord immunostained for ATF3 (grayscale top row, magenta in merge), Nissl (green), and DAPI (blue). ATF3 is a nuclear transcription factor; only nuclear signal is specific. Each knockout is validated against a positive control imaged in the same acquisition session. (A, B) Global knockout: in C57BL/6 (wild-type) DRG (A), nuclear ATF3 is present in a subset of sensory neurons; the identical display window applied to Atf3−/− (global-null) DRG from the same session (B) shows no nuclear ATF3. (C, D) Conditional knockout (*Avil*^Cre/+^; *Atf3*^fl/fl^): Advillin-Cre deletes Atf3 only in peripheral sensory neurons; ventral-horn motor neurons, axotomized by the same sciatic transection, retain ATF3 and serve as the internal positive control. The window that shows bright nuclear ATF3⁺ motor neurons in the conditional-knockout ventral spinal cord (C), applied identically to the same animal’s DRG (D), shows no nuclear ATF3, confirming a sensory-neuron-specific deletion. This validation is qualitative and confirms protein-level deletion; it is not a quantified measure of deletion efficiency. Scale bar, 100 µm.

**Supplementary Figure S3.**
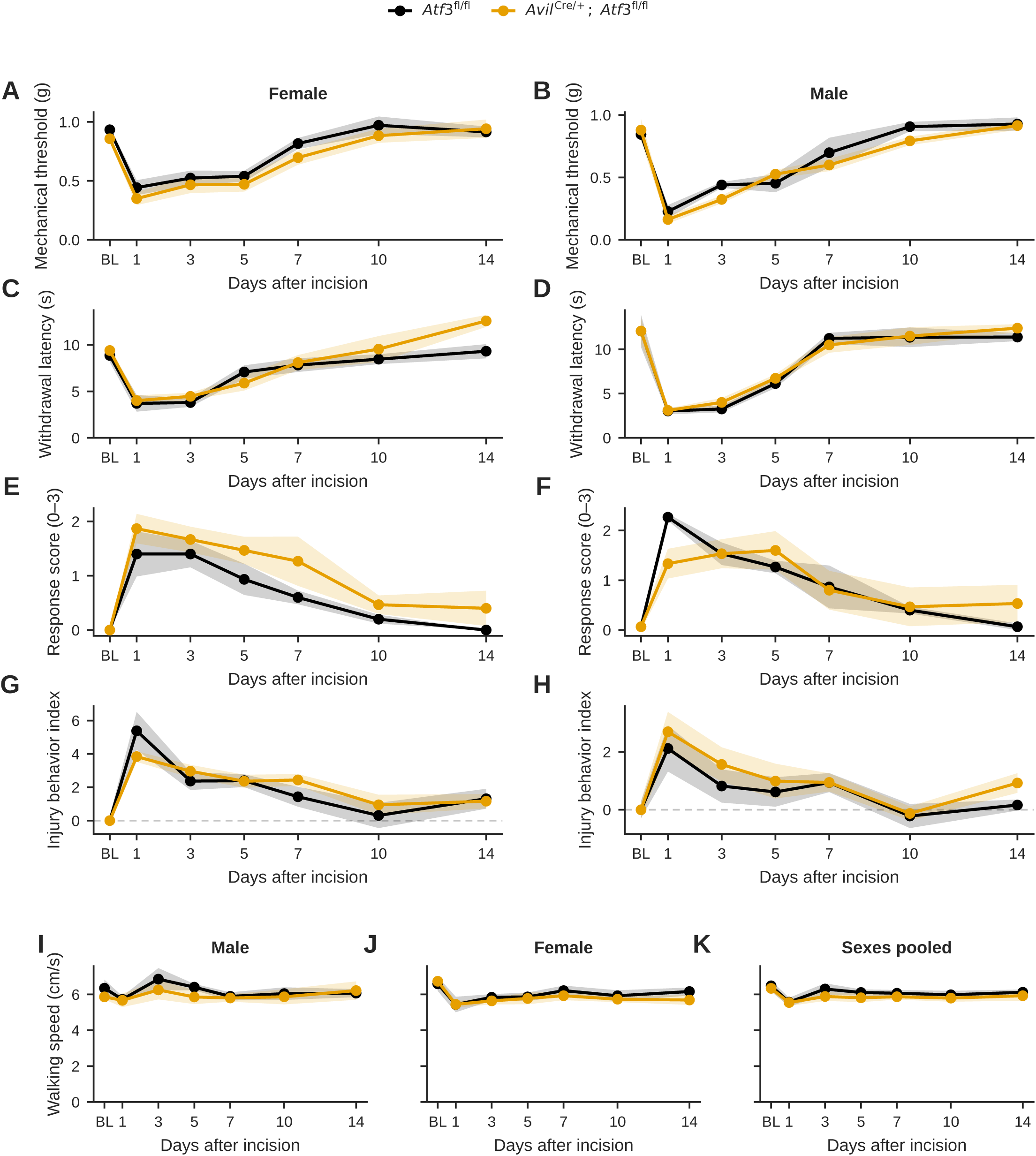
ATF3 deletion does not alter nociceptive behavior in either sex and does not impair locomotion. Behavioral support for Figure 2 (sensory-neuron Atf3 conditional knockout, *Avil*^Cre/+^; *Atf3*^fl/fl^ vs *Atf3*^fl/fl^ littermate control). Model: two-way mixed repeated-measures ANOVA (between = genotype, within = postoperative day; Greenhouse–Geisser corrected where sphericity was violated). (A–H) Behavior by sex: von Frey mechanical threshold (A, female; B, male), Hargreaves thermal latency (C, female; D, male), dynamic light-touch response score (E, female; F, male), and Black Box injury-behavior index (G, female; H, male); n = 5 per genotype per sex for the reflex assays (A–F), n = 6 (female)/n = 5 (male) per genotype for the injury index (G, H). The pooled genotype equivalence (Figure 2) holds within each sex: neither the genotype main effect nor the genotype × day interaction reached significance in any sex-stratified assay (all genotype P ≥ 0.12; all interaction P ≥ 0.19). (I–K) Locomotion: voluntary walking speed (cm/s) in male (I), female (J), and sex-pooled (K) cohorts (n = 5 male, 6 female, 11 pooled per genotype). Walking speed was indistinguishable between genotypes at every time point (pooled genotype F(1,19) = 0.81, P = 0.38; genotype × day F(6,114) = 0.23, P = 0.97) and poolable across sex (sex × genotype P = 0.76), supporting that the injury-behavior index reflects injury-related guarding rather than a generalized motor deficit. A modest parallel baseline-to-early-postoperative decline in both genotypes reflects arena habituation (main effect of day, P = 0.004).

**Supplementary Figure S4.**
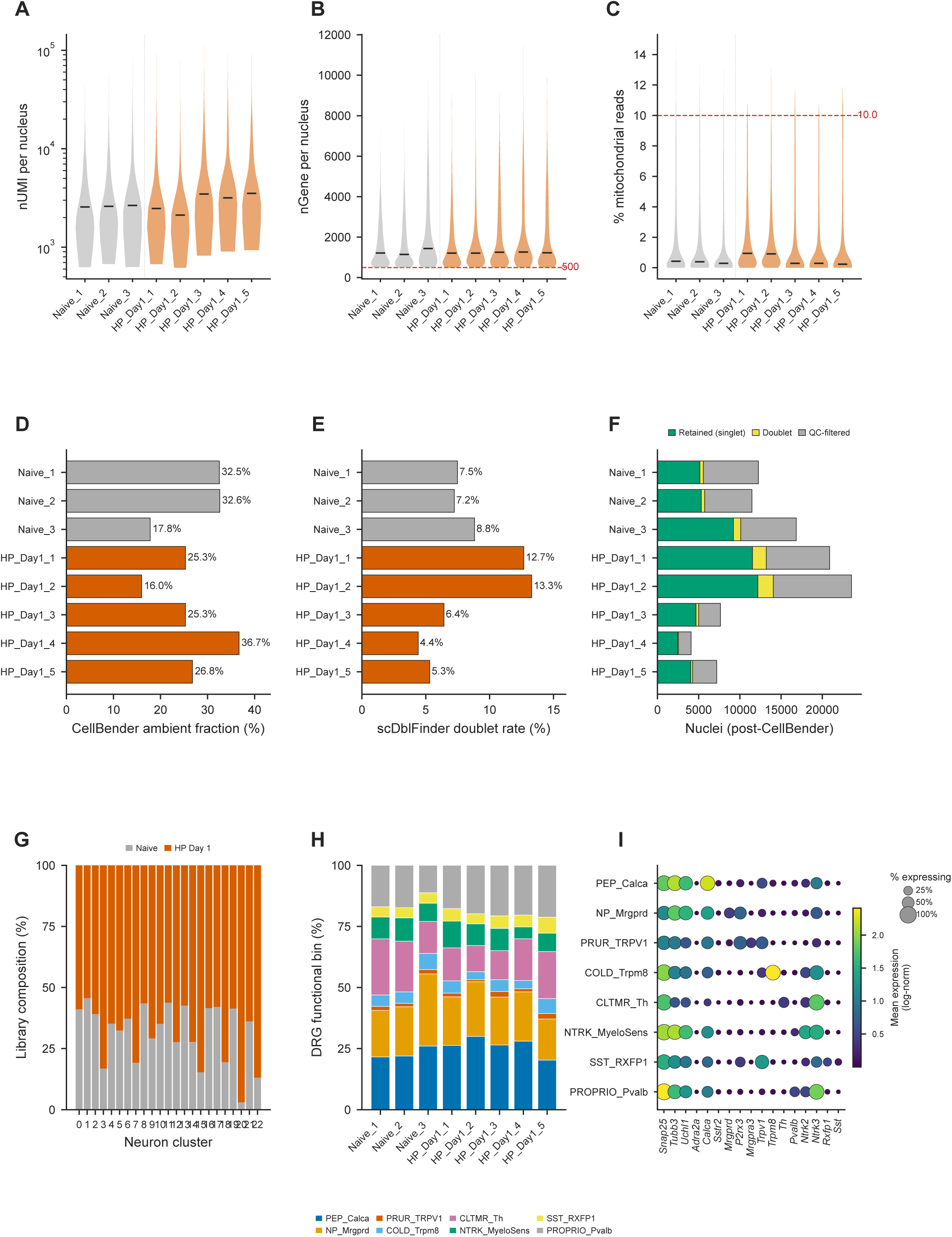
Single-nucleus RNA-sequencing quality control, wild-type (C57BL/6) cohort. Support for Figure 3. Per-sample QC for the wild-type snRNA-seq cohort (3 naïve, 5 hind-paw POD1). Processing: CellBender ambient-RNA removal, per-nucleus filtering (nGene ≥ 500, mitochondrial reads < 10%), scDblFinder doublet removal, and clustering with marker-based annotation. Dashed red lines mark filtering thresholds. (A) Total UMIs per nucleus (log scale), (B) genes detected per nucleus (threshold 500), and (C) percent mitochondrial reads (threshold 10%), by sample; black bars are medians. (D) CellBender-estimated ambient-RNA fraction per library. (E) scDblFinder doublet rate per library. (F) Nuclei per library after CellBender, partitioned into retained singlets, called doublets, and QC-filtered. (G) Per-cluster library composition (naïve vs POD1), confirming every cluster draws from both conditions. (H) Proportion of the eight annotated DRG neuronal subtypes per sample. (I) Canonical-marker dot plot across the eight subtypes (dot size = percent expressing; color = mean log-normalized expression), supporting the annotation.

**Supplementary Figure S5.**
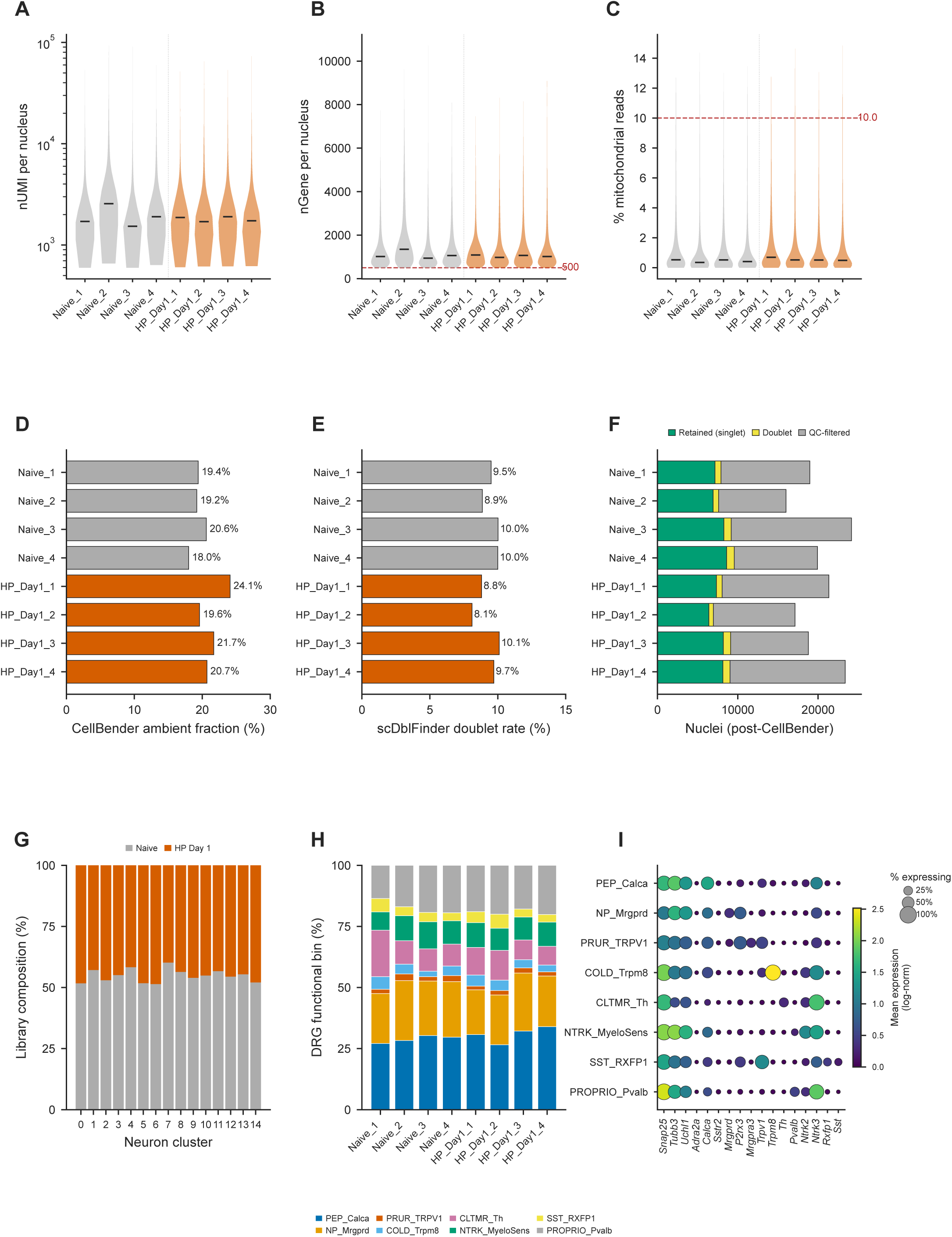
Single-nucleus RNA-sequencing quality control, Atf3-knockout cohort. As in Supplementary Figure S4, for the Atf3-knockout cohort (4 naïve, 4 hind-paw POD1), an identical processing pipeline, thresholds, and panel layout (A–I). QC metrics are comparable to those of the wild-type cohort, indicating that cross-genotype comparisons (Figures 3–4; Supplementary Figs. S6–S8) are not confounded by differences in library quality, ambient contamination, doublet rate, or capture depth between genotypes.

**Supplementary Figure S6.**
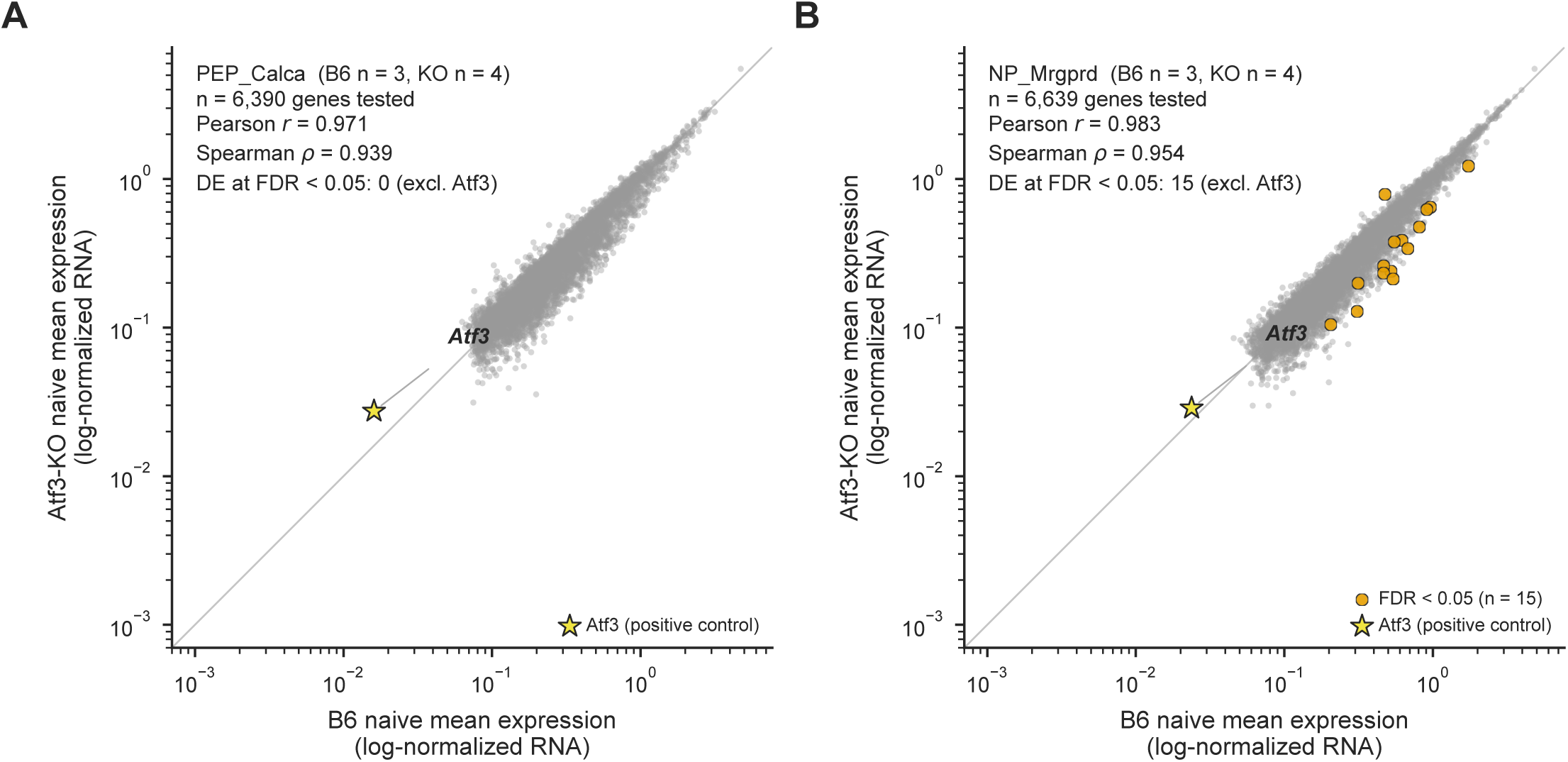
Naïve wild-type and Atf3-knockout sensory neurons are transcriptionally equivalent at baseline. Support for Figure 4: excludes a constitutive (developmental) transcriptional difference between genotypes as an explanation for the injury-response divergence. Per-gene mean expression (pseudobulk, log scale) is compared between naïve wild-type and naïve Atf3-knockout nuclei within each nociceptor lineage; each point is one gene, and the dashed line is the identity diagonal. *Atf3* itself sits off-diagonal as the expected positive control and is excluded from the concordance statistics. (**A**) Peptidergic (PEP_Calca) lineage: highly concordant (Pearson *r* = 0.97, Spearman ρ = 0.94); no gene other than *Atf3* is differentially expressed at baseline (false-discovery rate < 0.05). (**B**) Non-peptidergic (NP_Mrgprd) lineage: likewise concordant (Pearson *r* = 0.98, Spearman ρ = 0.95); aside from *Atf3*, only 15 genes reach baseline differential expression (false-discovery rate < 0.05), none belonging to the surgery-responsive functional categories tested in the main figures. The genotypes, therefore, start from an equivalent transcriptional baseline, so the divergent injury responses in Figures 3–4 reflect the incision response, not a developmental difference.

**Supplementary Figure S7.**
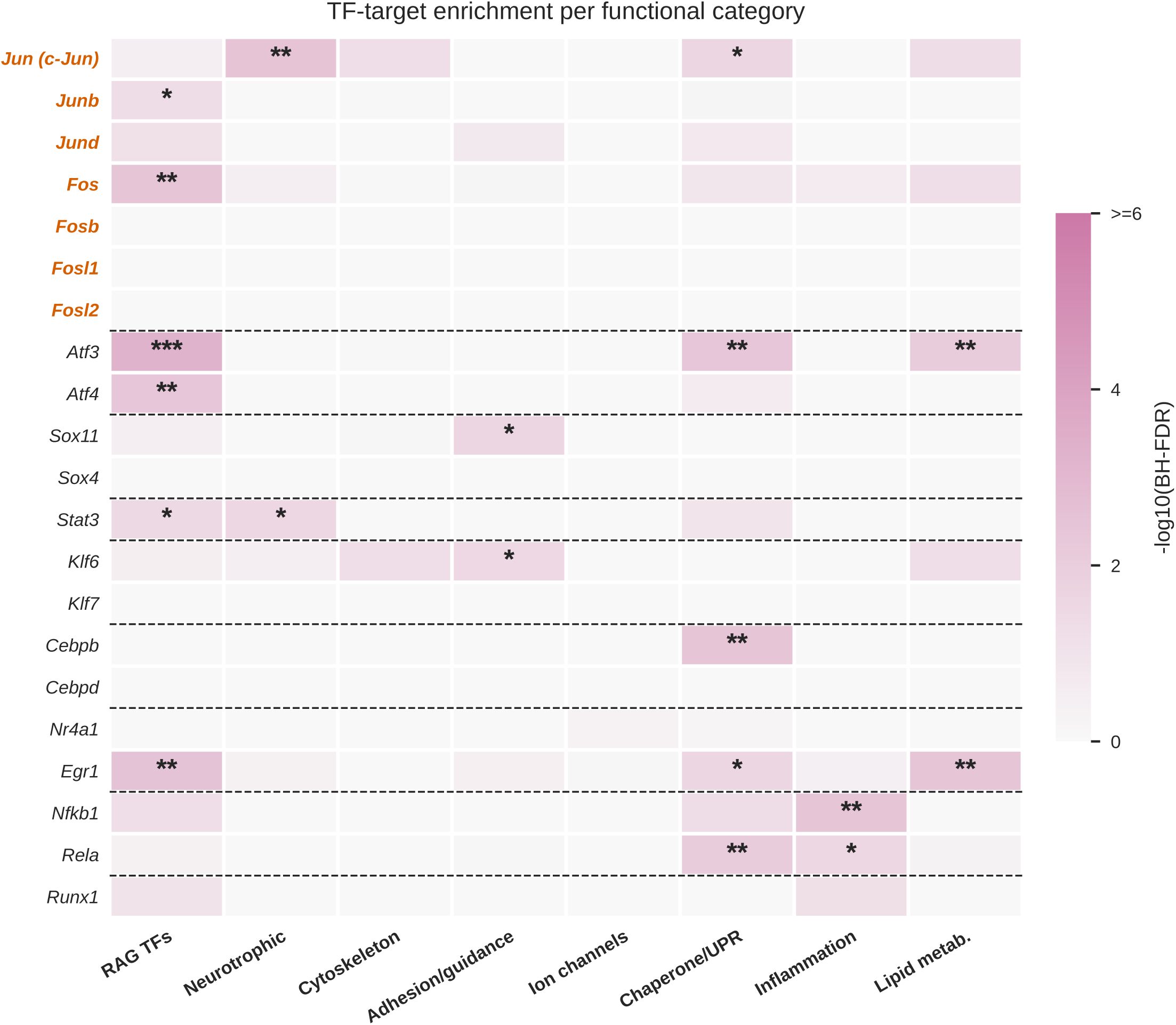
Transcription-factor target-set enrichment across the injury-program functional categories. Support for Figure 4C. An orthogonal, regulon-overlap test of which transcription factors regulate each of the eight injury-program functional categories, complementing the decoupleR activity inference in Figure 4C. For every transcription factor × category pair, over-representation of the factor’s target set among the category’s genes was assessed by one-sided Fisher’s exact test with Benjamini–Hochberg correction. Target sets are DoRothEA (A/B/C confidence) regulons; the AP-1 family is grouped at the top. Cell shading is −log₁₀(BH-FDR); stars mark significance. AP-1/Jun-family and other regeneration-associated factors are enriched across categories: c-Jun (*Jun*) targets are over-represented in the neurotrophic (BH-FDR = 3.6 × 10⁻^3^, 31-fold) and chaperone/UPR (BH-FDR = 0.025) categories; *Atf3* in regeneration-associated-transcription-factor (BH-FDR = 6.1 × 10⁻^4^), chaperone/UPR, and lipid categories; and *Atf4*, *Fos*, *Junb*, *Stat3*, and *Egr1* in the regeneration-associated-transcription-factor category. Inflammation-category genes are enriched for NF-κB-family (*Nfkb1*, *Rela*) targets. This target-set evidence corroborates the Figure 4C nomination of AP-1/c-Jun as an ATF3-independent factor associated with the injury program. *Sox11* and *Klf6* targets are enriched in the adhesion/axon-guidance category; *Sox11* is shown as one of the cooperative regeneration-associated factors in the panel and is not interpreted as a downstream ATF3 target.

**Supplementary Figure S8.**
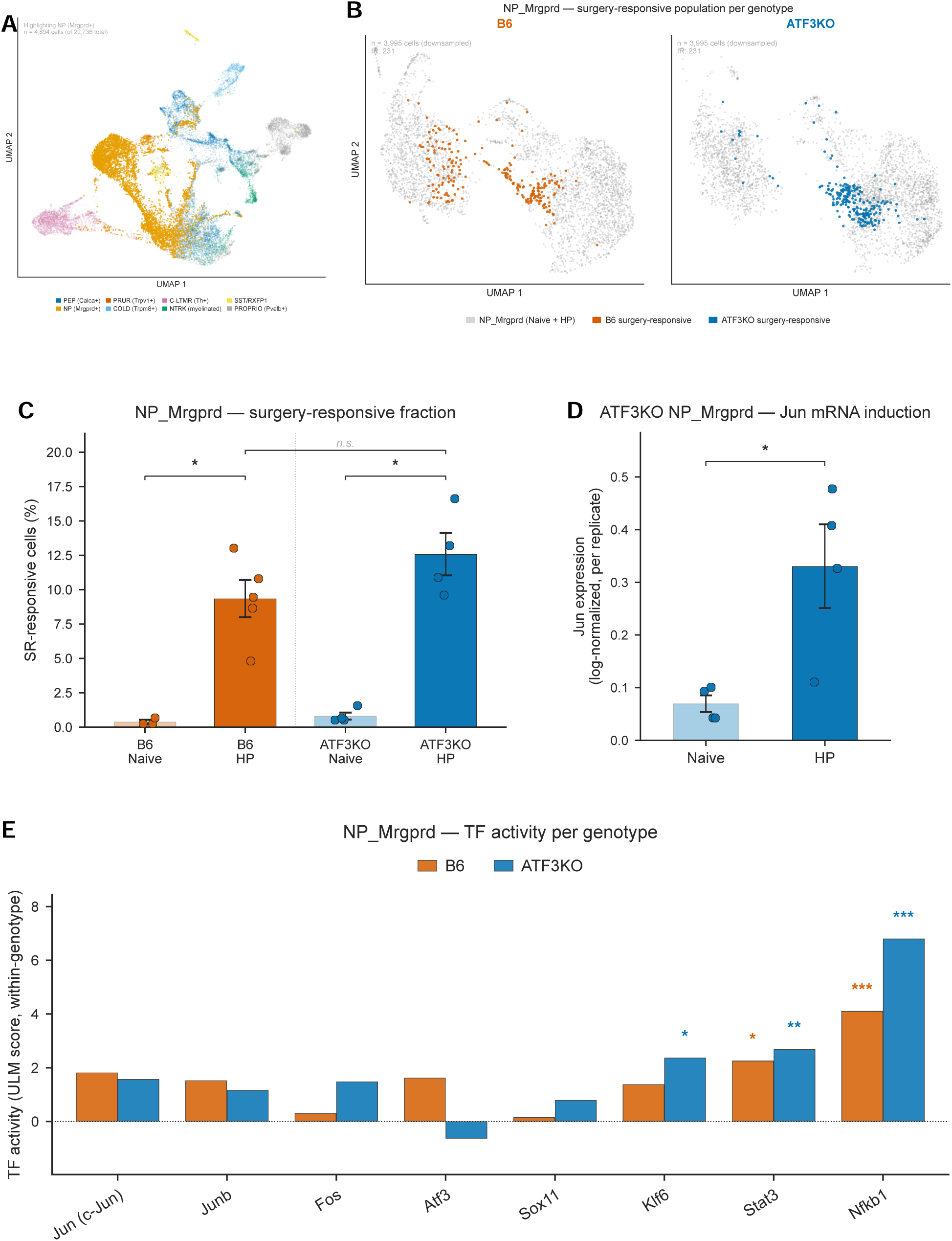
NP_Mrgprd cross-subtype sensitivity test for the cross-genotype findings. Support for Figures 3–4. The non-peptidergic (NP_Mrgprd, Mrgprd⁺) nociceptor lineage is a mechanistically distinct nociceptor class used to test the generality (not to provide a fully powered parallel analysis) of the PEP_Calca conclusions. (A) Reference UMAP of the full DRG snRNA-seq atlas (22,736 nuclei) with the NP_Mrgprd lineage highlighted (n = 4,894 nuclei). (B) NP_Mrgprd nuclei split by genotype (each downsampled to n = 3,995 for visual parity); surgery-responsive nuclei highlighted per genotype (SR = 231 each). (C) Per-replicate percentage of NP_Mrgprd singlets classified as surgery-responsive across the four groups (B6 naïve n = 3, B6 hind-paw n = 5, Atf3-knockout naïve n = 4, Atf3-knockout hind-paw n = 4); both genotypes mount a comparable injury response (within-genotype hind-paw vs naïve, two-sided Mann–Whitney U; between-genotype n.s.). (D) Atf3-knockout NP_Mrgprd per-replicate Jun mRNA (naïve n = 4 versus hind-paw surgery-responsive n = 4); Jun is induced by incision within the knockout (two-sided Wilcoxon rank-sum, P < 0.05), reproducing the peptidergic finding in a second nociceptor class. (E) Per-genotype transcription-factor activity (decoupleR) for the NP_Mrgprd lineage: AP-1/Jun-family activity is preserved across genotypes. As a descriptive, subtype-specific observation, Nfkb1 (NF-κB) activity is comparatively higher in the Atf3-knockout NP lineage; the NP cohort is underpowered, and this figure is a sensitivity check, not a parallel full analysis.

